# Experimental Validation of Finite Element Models for Directional DBS: The Critical Role of Boundary Conditions on VTA Accuracy

**DOI:** 10.64898/2026.04.03.716362

**Authors:** Kaylee R. Henry, Fuchang Jiang, William A. Wartman, Dexuan Tang, Yifeng Qian, Behzad Elahi, Sergey N. Makaroff, Laleh Golestanirad

## Abstract

**Objective:** Computational models and visualization toolboxes for Deep Brain Stimulation (DBS) increasingly rely on pre-computed electric field libraries to estimate the Volume of Tissue Activated (VTA). However, the boundary conditions (BCs) and source models used to generate these fields vary widely across studies, and there is currently no experimental consensus regarding which parameters most accurately reflect the physical device output. The objective of this study was to experimentally validate the electric potential distribution of directional DBS leads in order to determine the optimal Finite Element Method (FEM) configuration.

**Approach:** The voltage distribution surrounding a Boston Scientific Vercise Gevia directional lead was mapped in a saline phantom using a custom high-precision robotic scanning system. Experimental measurements were compared against six FEM configurations that varied in source formulation (Dirichlet vs. Neumann boundary conditions) and ground definitions. For each configuration, the resulting VTA volume was computed to assess the clinical impact of modeling assumptions.

**Results:** The FEM configuration implementing a Dirichlet (voltage) boundary condition on the active contact with a grounded implantable pulse generator (IPG) surface demonstrated the highest accuracy, achieving a Symmetric Mean Absolute Percent Error (SMAPE) of less than 9% across all contact levels. In contrast, conventional current-controlled simulations employing Neumann boundary conditions with disparate ground definitions substantially overestimated electric field spread. Suboptimal boundary condition selection resulted in an approximate 67% overestimation of VTA volume (137 mm^3^ vs. 82 mm^3^) relative to the experimentally validated model.

**Significance:** Although clinical DBS systems operate as current sources, standard Neumann (current density) boundary conditions do not adequately represent the equipotential behavior of the electrode–tissue interface, resulting in nearly a two-fold error in predicted VTA volume. To improve the validity of predictive clinical models, we recommend the use of Dirichlet boundary conditions derived from the device operating impedance (*V* = *I*_target_ × *Z*_measured_) rather than conventional current density specifications.

## 1 Introduction

Deep Brain Stimulation (DBS) is a critical therapeutic intervention for movement disorders, yet the clinical efficacy of the therapy is inextricably linked to the precise selection of stimulation parameters [1, 2, 3, 4]. The recent commercialization of directional leads with segmented contacts represents a significant hardware advance, offering the theoretical ability to “steer” the Volume of Tissue Activated (VTA) toward therapeutic targets and away from side-effect regions [5, 6, 7, 8]. However, this geometric flexibility introduces a “combinatorial explosion” in the programming parameter space. Clinicians are now faced with thousands of possible combinations of active segments, amplitudes, and current-steering ratios, making traditional trial-and-error programming effectively intractable [9].

To address this programming bottleneck, the field has increasingly pivoted toward computational solutions. A proliferation of automated optimization algorithms and visualization toolboxes aims to guide clinical decision-making by predicting the VTA for a given setting. The vast majority of these modern toolboxes, however, rely on a pre-computed field library—a static collection of electric potentials solved via finite element method (FEM) simulations for each individual electrode contact [10, 11, 12, 13]. Using the principle of superposition, these algorithms can instantaneously reconstruct the electric field for complex stimulation montages with negligible computational cost [13, 14]. Consequently, the validity of these high-level clinical tools rests entirely on the accuracy of the underlying, precomputed physics simulations. If the foundational electric field models are flawed, the downstream optimization recommendations will be systematically biased.

Despite this critical reliance, the parameters used to generate these simulations remain unstandardized and, crucially, unvalidated against experimental ground truth. The electric field is highly sensitive to multiple factors, including electrode geometry, tissue properties, and stimulation characteristics [15, 16]. While simulations must account for these variables, clinical information is often limited to the stimulation amplitude and the device-registered electrode-tissue impedance, leaving key modeling details—such as the specific stimulation type (current-controlled vs. voltage-controlled) and boundary assignments—to the user.

Modern clinical DBS implantable pulse generators (IPGs) are typically current-controlled devices, leading some researchers to model the active contact using a Neumann boundary condition [17, 18, 19], which enforces a specified current density flux across the electrode face. However, this approach introduces a conflict with material physics: highly conductive platinum-iridium contacts behave as equipotential surfaces. Enforcing a uniform current density in an FEM solver artificially forces the voltage to vary across the metal face, violating the equipotential constraint and potentially distorting the resulting field.

An alternative approach—applying a Dirichlet (voltage) boundary condition—respects the equipotential nature of the metal but requires the researcher to infer the voltage from device impedance. This methodological uncertainty has contributed to significant inconsistencies in the literature. Additionally, the location of the ground in the simulation is also up to interpretation. Some groups model the IPG casing or lower portion of the head and neck as the ground reference [17, 20], while others assign ground to the outer boundaries of the tissue domain [19, 21, 22]. Furthermore, a subset of studies does not report boundary definitions at all. Because the shape and volume of the VTA are highly sensitive to these potential gradients, these inconsistent modeling choices may have profound implications for clinical predictions.

In this study, we sought to resolve this ambiguity by establishing an experimental ground truth for the electric potential distribution generated by directional DBS leads. We constructed a custom, high-precision Cartesian robotic scanner to map the voltage field of a commercial Boston Scientific Vercise Gevia system submerged in a saline phantom. By comparing these experimental measurements against six distinct FEM configurations—varying the source type (voltage vs. current) and ground definition—we systematically evaluated which modeling strategy most faithfully reproduced physical reality. Our results provide a validated framework for defining simulation boundary conditions, helping ensure that future DBS optimization toolboxes are built upon a physically accurate foundation.

## 2 Methods

### 2.1 Experimental Measurement Setup

To validate the electric potential distribution, we designed a high-precision robotic scanning platform to map the voltage field generated by a commercial directional DBS system in a saline phantom. A schematic overview of the system architecture is provided in Fig. 1(a), and photographs of the physical setup are available in Supplementary Fig. 1.

**Figure 1.**
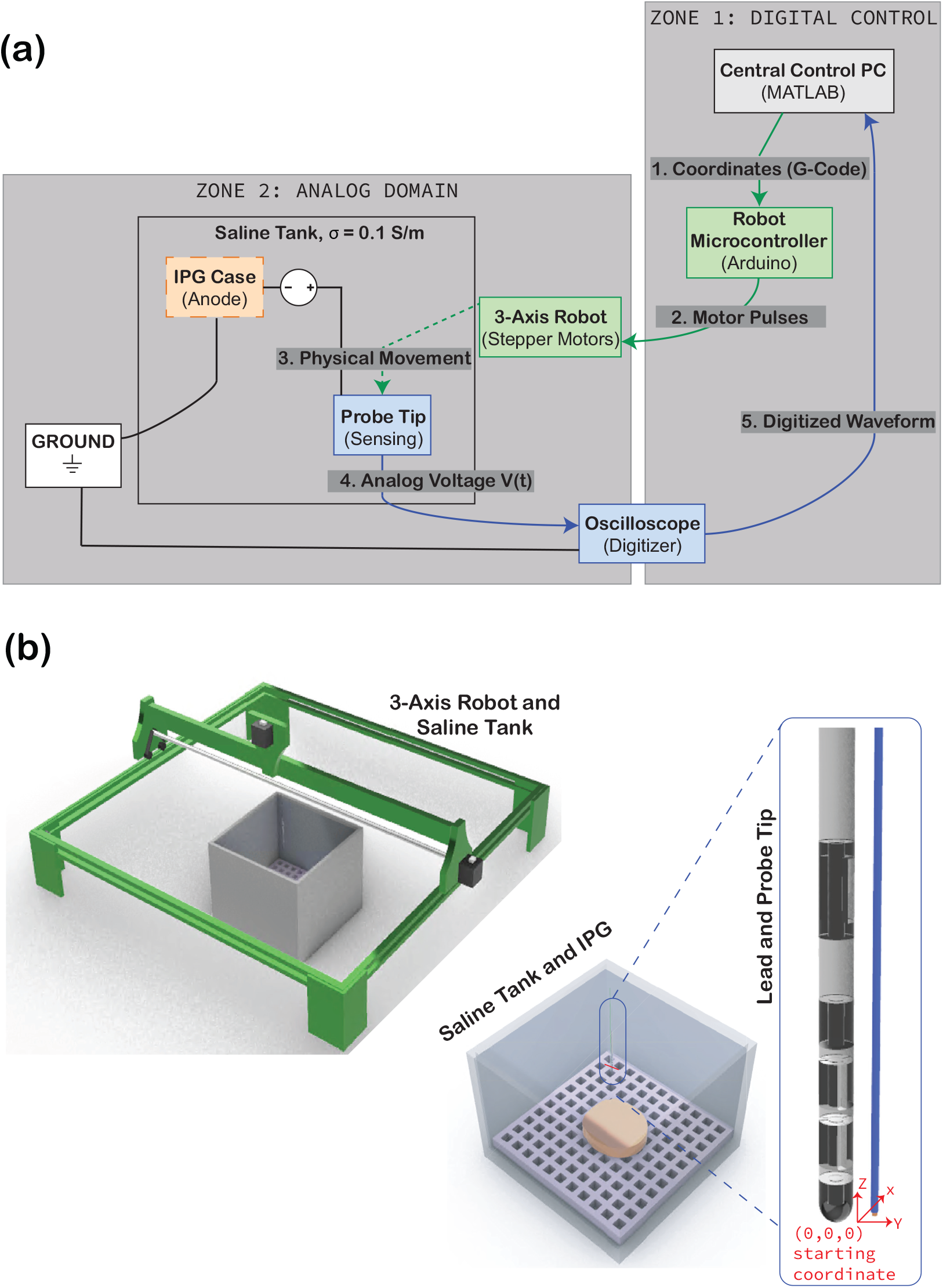
High-precision experimental validation platform. (a) System architecture and signal flow. The Central Control PC synchronizes the motion control loop (green path: PC → Robot Microcontroller → Robot) and the data acquisition loop (blue path: Probe → Oscilloscope → PC). (b) Three-dimensional spatial implementation. CAD rendering of the custom Cartesian robot and saline phantom. Components are color-coded to correspond with the schematic: the mechanical gantry (green) positions the sensing element, while the submerged IPG and probe (blue) form the measurement circuit.

#### 2.2.1 Custom Cartesian Robot and Device Holder

A 3-axis Cartesian coordinate robot was constructed by modifying a Sculpfun S9 laser engraver frame (Shenzhen Sculpfun Technology Co., Guangdong, China). The motion system utilized NEMA-17 stepper motors driven by an Arduino Mega (Arduino, Somerville, MA, USA) equipped with a RAMPS 1.4 shield running Marlin firmware (https://marlinfw.org/). Positioning was controlled via G-code commands sent from MATLAB 2023a (The MathWorks, Natick, MA, USA), achieving a spatial step size of 0.2 mm. The motion system was powered by an external RIGOL DP832A programmable 12 V DC power supply (RIGOL Technologies, Beijing, China).

A custom DBS device holder (Fig. 2(a)) was designed in Rhinoceros 3D (Robert McNeel & Associates, Seattle, WA, USA) to rigidly secure the IPG and lead in a fixed geometry. The assembly consisted of three components: a main container (15 × 15× 11 cm) with a securing grid, a form-fitting IPG holder, and a lead holder. To ensure mechanical stability and dielectric compatibility, the container and IPG holder were fabricated using Tough 1500 Resin on a Formlabs SLA printer (Formlabs Inc., Somerville, MA, USA). The lead holder was designed to hold the lead upright, directly centered over the IPG at the center of the container and was printed in PLA using Fused Deposition Modeling (FDM) 3D printer (Bambu Lab, Shenzhen, China).

**Figure 2.**
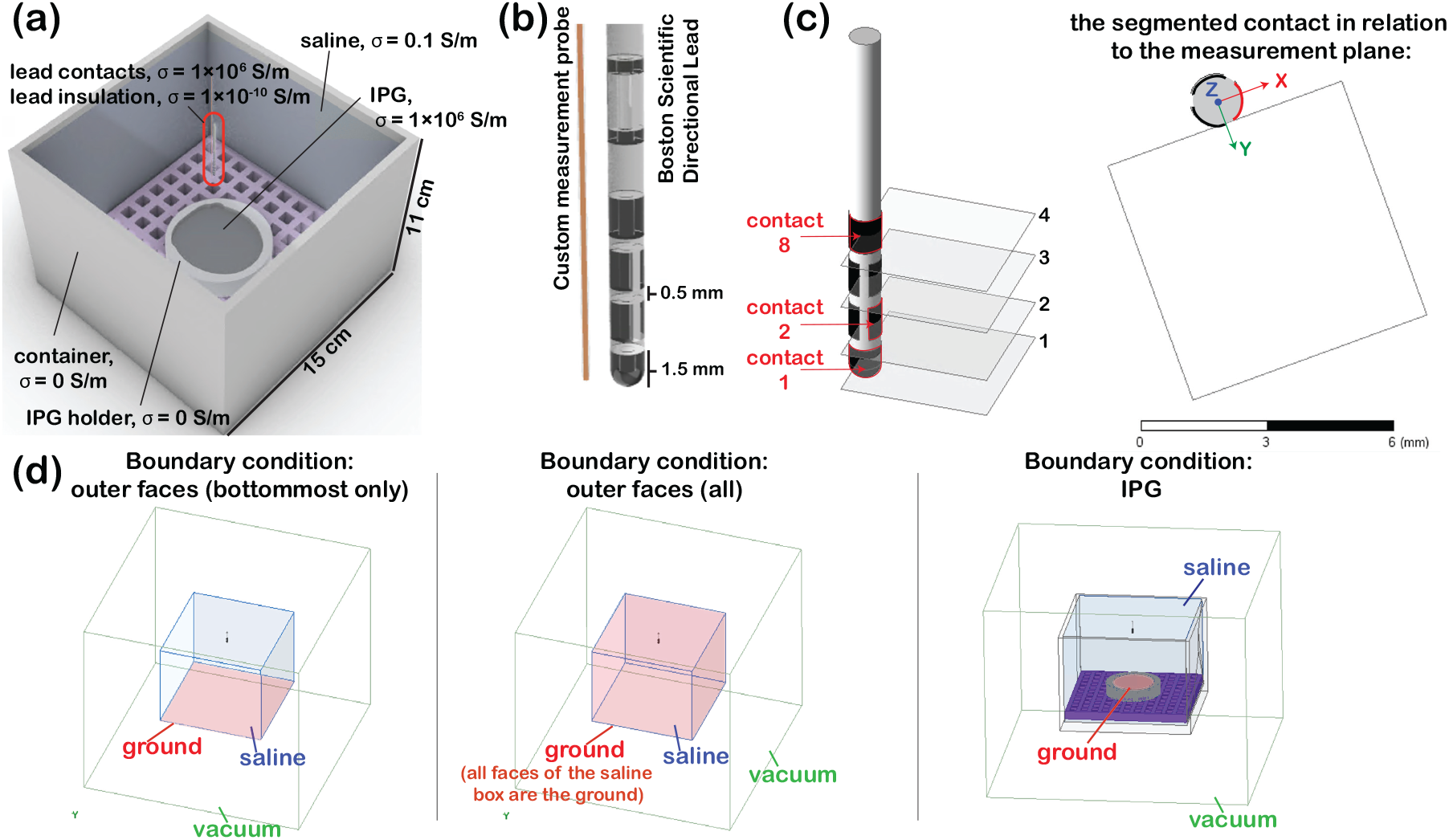
Finite Element Model (FEM) parameters. (a) Geometry and conductivity values assigned to the lead, insulation, saline, and IPG. (b) Detail of the directional lead geometry and the custom measurement probe. (c) Location of the four horizontal 6 mm × 6 mm measurement planes (Levels 1–4) in relation to contacts 1, 2, and 8. (d) Visualization of the three ground boundary condition strategies tested. Red surfaces indicate the application of the 0 V Dirichlet boundary condition.

The container was filled with a saline solution titrated to a conductivity of 0.1 S/m (measured using a SANXIN Model MP513 conductivity meter, Shanghai San-Xin Instrumentation, Inc.) to mimic the average bulk conductivity of intracranial tissue (Fig. 2(a)). A custom measurement probe was fabricated using 0.2 mm insulated magnet wire threaded through a 100 mm long, 0.3 mm diameter capillary tube (Fig. 2(b)). The insulation at the tip was removed via manual abrasion with sandpaper to expose the conductor.

#### 2.1.2 DBS Stimulation and Data Acquisition

We utilized a Boston Scientific Vercise Gevia system with a standard directional lead (1.5 mm contacts, 0.5 mm spacing). The device was programmed to a monopolar configuration (IPG case as anode, Contact 1 as cathode) with a regulated current amplitude of 5 mA. Directional stimulation was tested using Contact 2 as the cathode. Device impedance was recorded using the clinical programmer prior to each trial. Data were collected along four horizontal planes (Levels 1–4) corresponding to the vertical position of each contact on the lead Fig. 2(c)).

Voltage waveforms were acquired using a RIGOL DS2202A digital oscilloscope communicating with MATLAB via USB using a sampling rate of 1 GSa/s. To mitigate high-frequency noise, a custom one-dimensional moving-average filter (window length = 5 samples) was applied to the raw waveform (Fig. 3). This window size was validated to preserve peak voltage characteristics without attenuating the signal amplitude (Supplementary Fig. 2). Peak-to-peak voltages were extracted from the filtered signal at each spatial coordinate.

**Figure 3.**
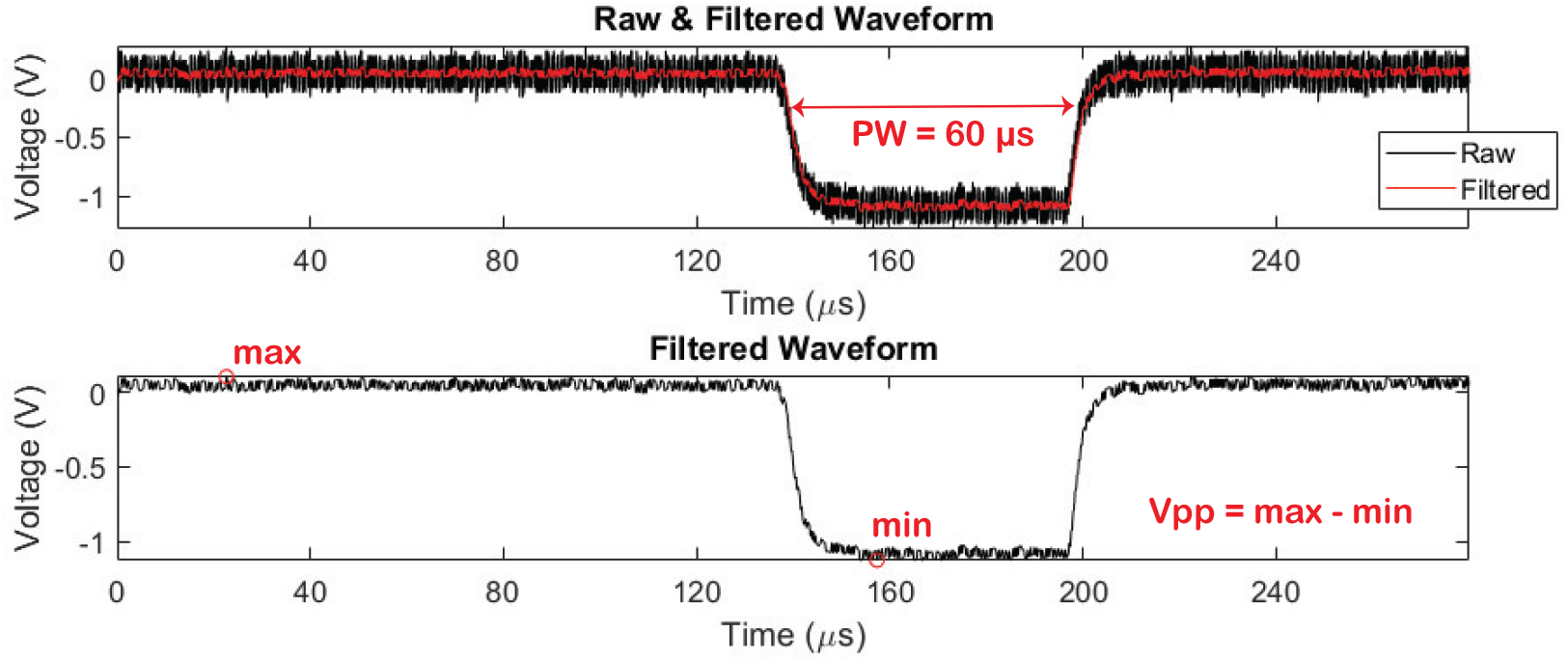
Signal processing pipeline. Top: Raw voltage waveform acquired from the oscilloscope (black) overlaid with the filtered signal (red), pulse width = 60 *µ*s. Bottom: The filtered waveform used to extract the peak-to-peak voltage (*V*_pp_). A moving-average filter with a window of 5 samples (approx. 1 *µ*s) was applied to mitigate high-frequency noise while preserving the pulse rise time and amplitude characteristics.

### 2.2 Finite Element Method (FEM) Simulations

3D FEM simulations were carried out using Maxwell in ANSYS Electronics Desktop 2023 R1 (Ansys Inc., Canonsburg, PA, USA).

#### 2.2.1 Governing Equations and Quasi-Static Assumption

Although clinical DBS employs pulsed stimulation, the associated electromagnetic wavelengths at therapeutic frequencies are several orders of magnitude larger than the dimensions of the volume conductor. Consequently, the quasi-static approximation (*σ* ≫ *ωε*) is valid, and the electric potential distribution *V* can be obtained by solving the Laplace equation,

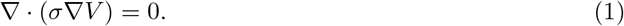

The saline domain was modeled with a conductivity of *σ* = 0.1 S/m. The implantable pulse generator (IPG) and lead contacts were modeled as nearly perfect conductors (*σ* = 1 × 10^6^ S/m), whereas the lead insulation and holder were treated as electrical insulators (*σ* ≈ 0 S/m), as shown in Fig. 2(a).

#### 2.2.2 Boundary Conditions and Source Models

We compared six simulation configurations varying in source type (Fig. 2(d)):

a. Neumann Condition (current-controlled): To model the device as a current source, a 5 mA current excitation was applied to the active contact (Contact 1 or 2). In standard FEM solvers, this enforces a uniform current density flux across the contact surface.
b. Dirichlet Condition (voltage-controlled): To model the equipotential nature of the electrode, a fixed voltage excitation was applied. The magnitude was derived from the experimentally measured impedance (*Z*) and target current (*I*) via Ohm’s law (*V* = *I* × *Z*). For the omnidirectional case (*Z* ≈ 1412 Ω), the excitation was set to −7.06 V. For the directional case (*Z* ≈ 2354 Ω) the excitation was set to −11.8 V.

Then, for each source type, we tested three ground (0 V Dirichlet) definitions: (i) IPG Surface: ground applied exclusively to the IPG surface; (ii) Outer Faces (All): ground applied to all outer faces of the saline container; (iii) Outer Faces (Bottommost): ground applied only to the bottom face of the container.

#### 2.2.3 Impedance Matching

To separate the effects of boundary conditions from conductivity mismatches, a second set of simulations was run where the saline conductivity was adjusted until the simulated impedance matched the clinical device impedance (± 5 Ω)

### 2.3 Volume of Tissue Activated (VTA) Estimation

To assess clinical impact, the activating function was approximated using the Hessian matrix (*H*) of the second spatial derivatives of the electric potential (*V*_*e*_): [11, 23, 24, 25].

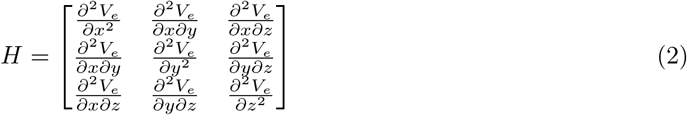

The Hessian was computed within a 20 × 20 × 20 mm volume (0.1 mm resolution) around the lead. Following the AF-3D method introduced by Duffley et al. [21], the maximum eigenvalue of *H* was computed at each point. A standard activation threshold of 26.7 V/cm^2^ was applied to define the VTA, representing the activation of myelinated axons (5.7 *µ*m diameter, 70 Ω cm resistivity [26]).

### 2.4 Statistical Analysis

Simulation accuracy was quantified using a variation of the Symmetric Mean Absolute Percent Error (SMAPE) bounded between 0% and 100%. We calculated the error at each spatial coordinate *t* as the absolute difference normalized by the sum of absolute values:

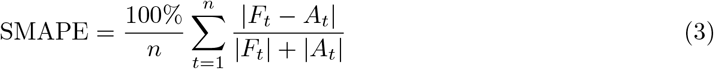

where *A*_*t*_ is the experimental voltage and *F*_*t*_ is the simulated voltage. Comparisons of absolute error distributions were performed using two-sample *t*-tests (*α* = 0.05) in MATLAB.

## 3 Results

### 3.1 Experimental Ground Truth

The robotic scanning platform successfully mapped the steady-state voltage distribution of the Boston Scientific Vercise Gevia system. The measured potentials exhibited the expected monotonic decay with distance. For omnidirectional stimulation (Level 1), voltages ranged from -6.89 V at the contact surface to -1.38 V at the periphery of the scan volume (Fig. 4, left column). This was also confirmed with an additional experiment where Contact 8 was used (Supp. Table 1). For directional stimulation (Level 2), the focused field resulted in a steeper gradient, with potentials ranging from -10.3 V to -0.98 V. These experimental datasets served as the ground truth for all subsequent simulation comparisons. All other simulation results are illustrated in Supplementary Figs. 3 and 4, and mesh details are available in Supplementary Tables 2-7.

**Table 1.** Comparison of simulated and experimental voltage distributions prior to impedance matching. The SMAPE (%) indicates the deviation between the model predictions and physical measurements. The Voltage-controlled (Dirichlet) setup with the grounded IPG case yields the lowest error. The reported Z values are the simulated values (i.e. we used the experimentally measured Z to calculate what V to apply, and the simulation resulted in a slightly different Z value, which is reported here).

| Simulation Setup | Level | Simulated<br>Min (V), Max (V) | Simulation vs.<br>Experiment: SMAPE (%) |
| --- | --- | --- | --- |
| Dirichlet, IPG<br>($\sigma = 0.1$ S/m, $Z_{\text{sim}} = 1206 \Omega$) | 1 | -7.06, -1.04 | 8.44 |
|  | 2 | -2.53, -1.02 | 1.49 |
|  | 3 | -1.42, -0.95 | 3.50 |
|  | 4 | -1.10, -0.87 | 4.11 |
| Neumann, IPG<br>($\sigma = 0.1$ S/m, $Z_{\text{sim}} = 1504 \Omega$) | 1 | -7.51, -0.89 | 14.87 |
|  | 2 | -2.39, -0.88 | 5.37 |
|  | 3 | -1.27, -0.82 | 5.94 |
|  | 4 | -0.97, -0.76 | 7.76 |
| Dirichlet, outer faces (all)<br>($\sigma = 0.1$ S/m, $Z_{\text{sim}} = 1127 \Omega$) | 1 | -7.06, -0.62 | 21.55 |
|  | 2 | -2.19, -0.58 | 15.89 |
|  | 3 | -0.99, -0.49 | 22.65 |
|  | 4 | -0.63, -0.40 | 31.94 |
| Neumann, outer faces (all)<br>($\sigma = 0.1$ S/m, $Z_{\text{sim}} = 2208 \Omega$) | 1 | -11.04, -0.98 | 7.29 |
|  | 2 | -3.47, -0.93 | 8.15 |
|  | 3 | -1.57, -0.78 | 6.76 |
|  | 4 | -1.00, -0.63 | 10.12 |
| Dirichlet, outer faces (bottommost only)<br>($\sigma = 0.1$ S/m, $Z_{\text{sim}} = 1175 \Omega$) | 1 | -7.06, -0.87 | 13.19 |
|  | 2 | -2.40, -0.85 | 5.16 |
|  | 3 | -1.26, -0.77 | 7.13 |
|  | 4 | -0.93, -0.70 | 10.45 |
| Neumann, outer faces (bottommost only)<br>($\sigma = 0.1$ S/m, $Z_{\text{sim}} = 2300 \Omega$) | 1 | -11.50, -1.45 | 12.08 |
|  | 2 | -3.96, -1.41 | 19.91 |
|  | 3 | -2.08, -1.29 | 18.09 |
|  | 4 | -1.54, -1.16 | 14.71 |

**Table 2.** Comparison of simulated and experimental voltage distributions after impedance matching. The table displays the updated SMAPE (%) alongside the original values (in parentheses) to illustrate model sensitivity. While impedance matching improved accuracy for specific Current-controlled (Neumann) configurations (e.g., bottommost ground), it drastically degraded others (e.g., all outer faces), highlighting the Neumann condition’s unpredictability and sensitivity to boundary assumptions. In contrast, the Voltage-controlled (Dirichlet) setup remained stable and robust across all conditions.

| Simulation Setup | Level | Simulated Min (V), Max (V) | Simulation vs. Experiment: SMAPE (%) (Original SMAPE Value from Table 1) |
| --- | --- | --- | --- |
| Dirichlet, IPG<br>($\sigma = 0.855$ S/m, $Z = 1412 \Omega$) | 1 | -7.06, -1.04 | 8.4 (8.4) |
|  | 2 | -2.53, -1.02 | 1.5 (1.5) |
|  | 3 | -1.42, -0.95 | 3.5 (3.5) |
|  | 4 | -1.10, -0.87 | 4.1 (4.1) |
| Neumann, IPG<br>($\sigma = 0.107$ S/m, $Z = 1412 \Omega$) | 1 | -7.06, -0.84 | 17.8 (14.9) |
|  | 2 | -2.25, -0.83 | 8.5 (5.4) |
|  | 3 | -1.20, -0.77 | 8.8 (5.9) |
|  | 4 | -0.91, -0.71 | 10.8 (7.8) |
| Dirichlet, outer faces (all)<br>($\sigma = 0.08$ S/m, $Z = 1412 \Omega$) | 1 | -7.06, -0.62 | 22.6 (21.6) |
|  | 2 | -2.19, -0.58 | 15.9 (15.9) |
|  | 3 | -0.99, -0.49 | 22.7 (22.7) |
|  | 4 | -0.63, -0.40 | 31.9 (31.9) |
| Neumann, outer faces (all)<br>($\sigma = 0.157$ S/m, $Z = 1414 \Omega$) | 1 | -7.03, -0.63 | 20.8 (7.3) |
|  | 2 | -2.21, -0.59 | 15.2 (8.2) |
|  | 3 | -1.00, -0.50 | 22.0 (6.8) |
|  | 4 | -0.64, -0.40 | 31.3 (10.1) |
| Dirichlet, outer faces<br>(bottommost only)<br>($\sigma = 0.083$ S/m, $Z = 1415 \Omega$) | 1 | -7.06, -0.87 | 13.2 (13.2) |
|  | 2 | -2.40, -0.85 | 5.1 (5.1) |
|  | 3 | -1.26, -0.77 | 7.1 (7.1) |
|  | 4 | -0.93, -0.70 | 10.4 (10.4) |
| Neumann, outer faces<br>(bottommost only)<br>($\sigma = 0.163$ S/m, $Z = 1411 \Omega$) | 1 | -7.05, -0.89 | 12.3 (12.1) |
|  | 2 | -2.43, -0.87 | 4.4 (19.9) |
|  | 3 | -1.28, -0.79 | 6.4 (18.1) |
|  | 4 | -0.94, -0.72 | 9.5 (14.7) |

**Figure 4.**
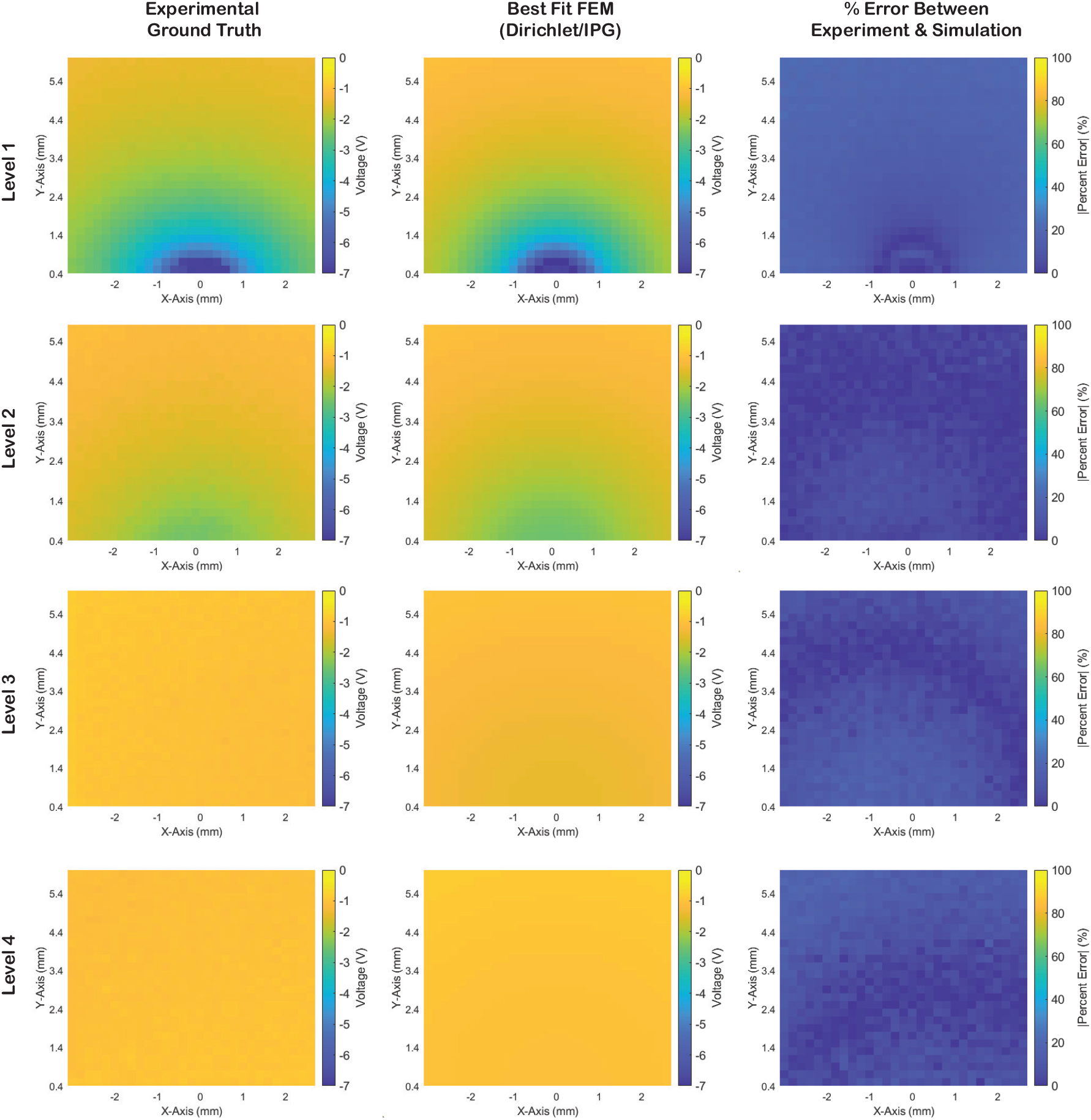
Validation of the optimal FEM configuration. Left: Experimental voltage distributions mapped by the robotic scanner at four vertical levels. Center: Simulated distributions using the optimal Dirichlet source with IPG ground. Right: Spatial error maps showing high agreement (SMAPE ¡ 9%) throughout the volume.

### 3.2 Identification of the Optimal Simulation Configuration

Comparison of the six FEM configurations revealed that the Dirichlet source with an IPG-surface ground provided the highest fidelity to the experimental data. This configuration achieved a Symmetric Mean Absolute Percent Error (SMAPE) of *<* 9% across all four vertical levels of the lead (Table 1). Specifically, the absolute percent error distribution for this model was significantly lower than competing configurations (*p <* 0.05), with interquartile ranges (IQR) of error remaining below 8% at all measured depths (Fig. 5). The VTA predicted by this validated model was 82.2 mm^3^.

**Figure 5.**
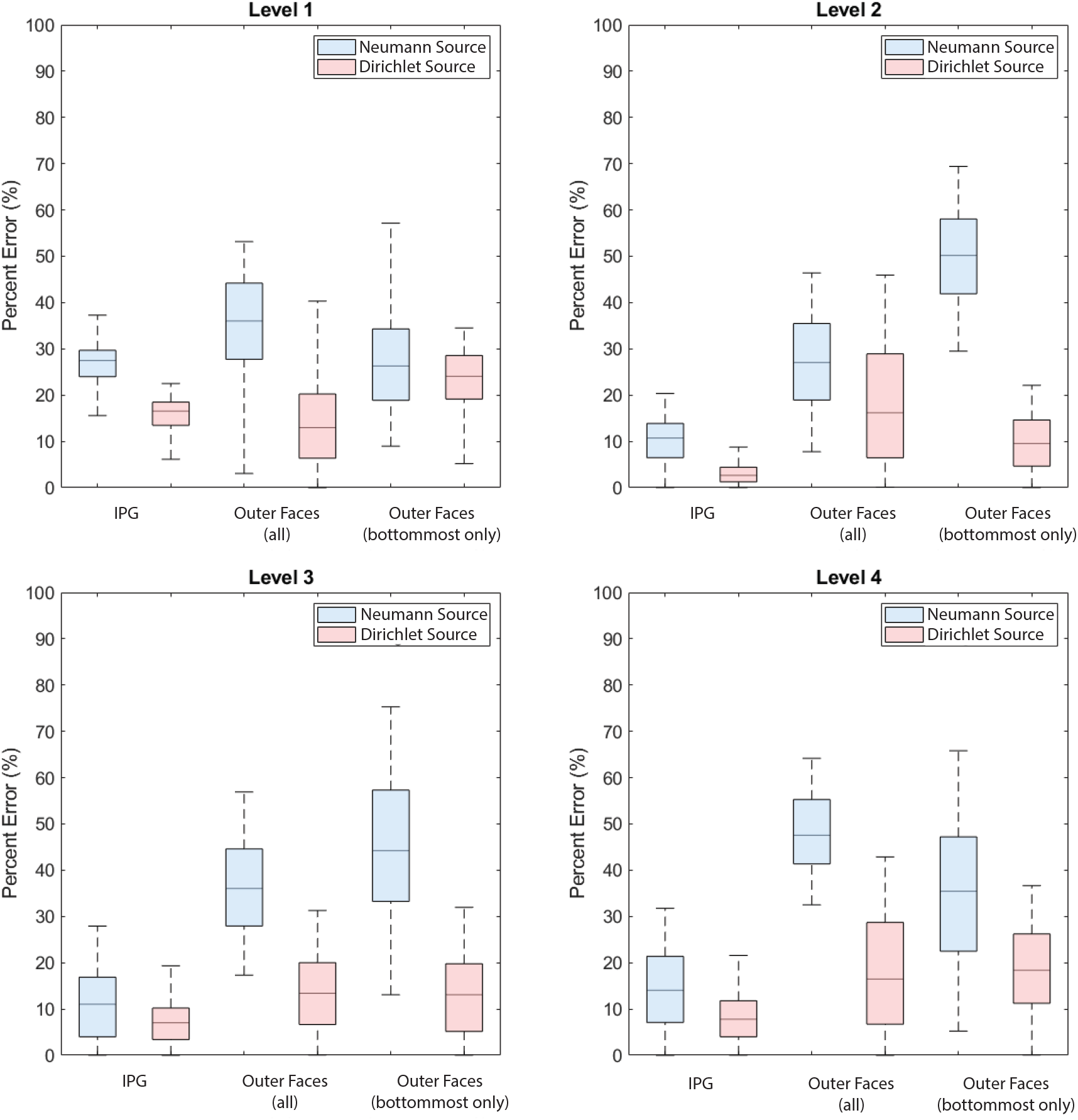
Quantitative accuracy of FEM configurations. Absolute percent error distributions comparing the experimental data to six simulation scenarios (varying source type and ground location) across all contact levels. The Dirichlet (Voltage-Controlled) source with IPG ground (red box, far left) consistently yields the lowest error distribution.

### 3.3 Consequences of Suboptimal Boundary Condition (VTA Overestimation)

Simulations utilizing standard Neumann boundary conditions with distant grounds—a common approach in the literature—resulted in substantial deviations from the experimental ground truth. Specifically, the Neumann model with “Outer Faces (All)” set as ground yielded significantly higher error rates, particularly in the high-gradient region near the contact interface (Fig. 5). Most critically, this choice of boundary condition dramatically altered the predicted clinical volume.

While the validated Dirichlet model predicted a VTA of ∼82 mm^3^, the Neumann model with the outer faces (all) grounds predicted a VTA of ∼137 mm^3^ (Fig. 6(a)). This represents an approximately 67% overestimation of neural activation volume solely due to the selection of boundary parameters.

**Figure 6.**
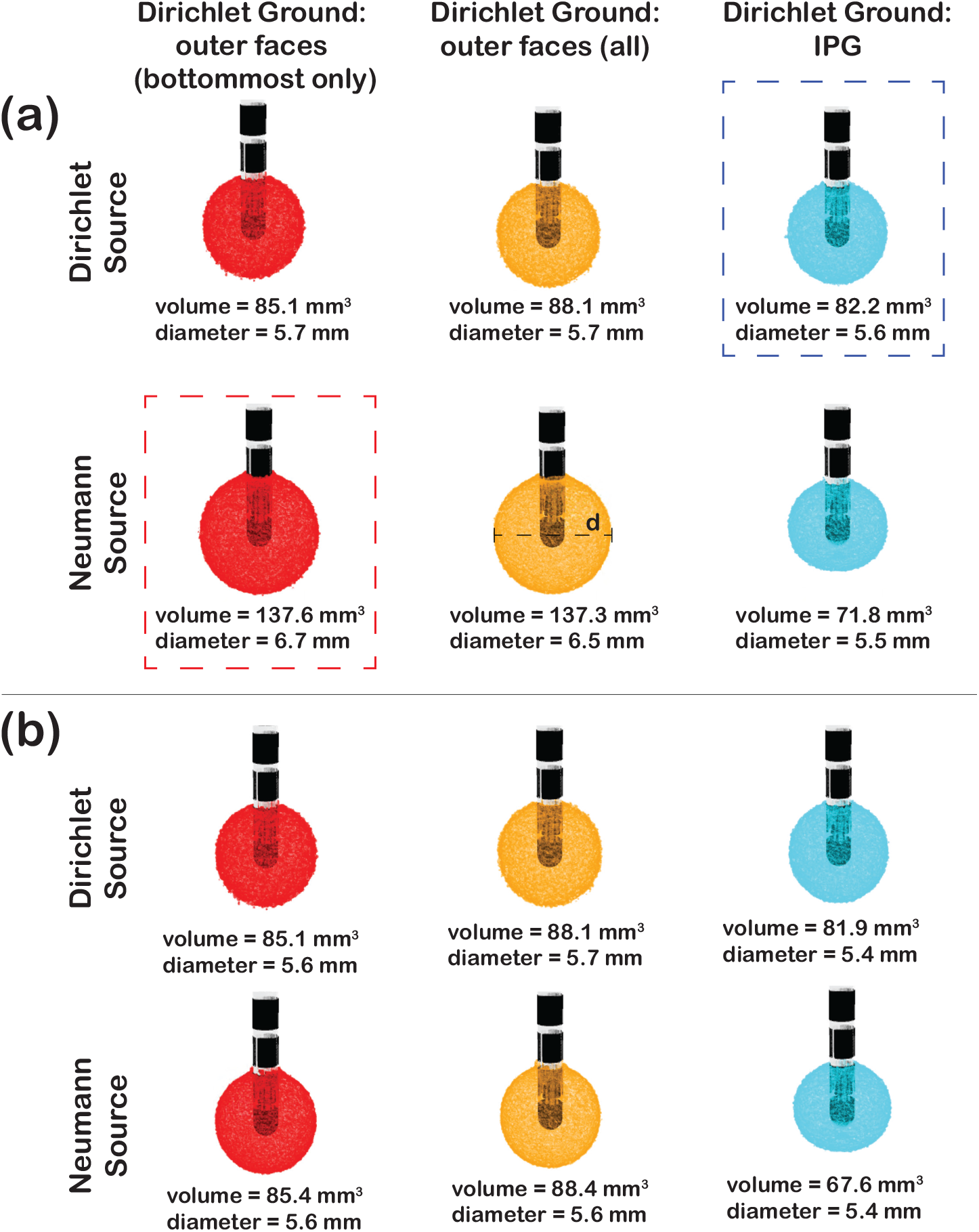
Clinical impact of boundary condition selection. (a) Comparison of VTAs generated by standard vs. optimized models. The standard Neumann Source (outlined in red dashed lines) and the best FEM (Dirichlet Source/IPG Ground) (outlined in blue dashed lines) are emphasized. (b) VTA predictions after matching simulation impedance to clinical impedance.

### 3.4 Impact of Impedance Matching

To determine if conductivity mismatches drove these errors, we performed a secondary analysis where the saline conductivity in each simulation was iteratively adjusted until the modelled impedance matched the clinical device impedance (± 5 Ω) (Table 2). Even after impedance matching, the Dirichlet configuration with IPG ground remained the most accurate, maintaining SMAPE values *<* 9% (Fig. 7). However, impedance matching did improve the performance of the Neumann models (Supplementary Fig. 5 and 6). Notably, the Neumann simulation with a “Bottommost Face” ground improved, achieving SMAPE values *<* 12% across all levels (Table 2). This suggests that if a Neumann solver is required, constraining the ground path to the bottommost surface (simulating a distal return) and strictly matching impedance is the second-best approximation.

**Figure 7.**
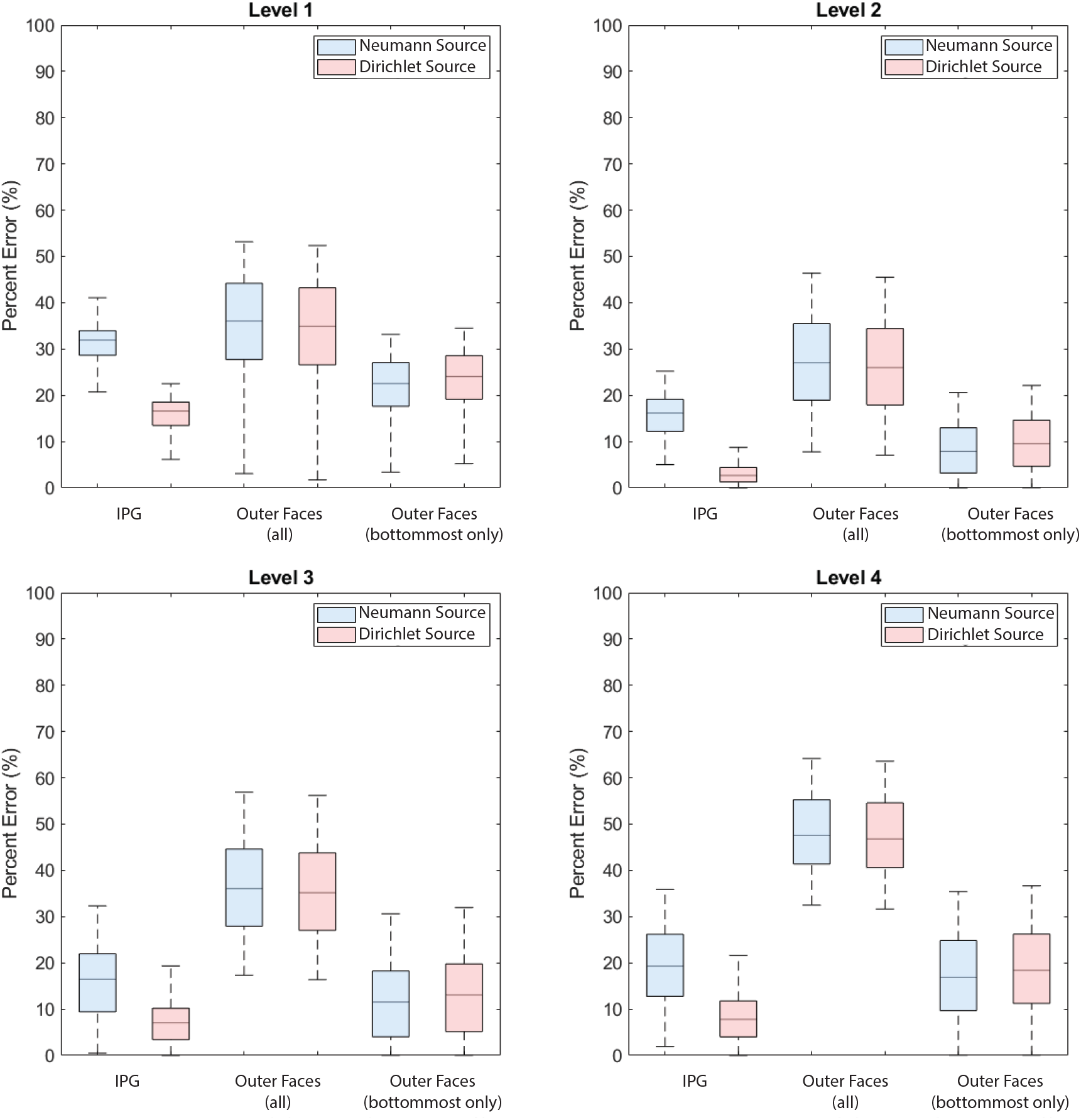
The absolute percent error between the experimental voltage distributions and simulated voltage distributions at each coordinate point on each level of contacts after matching the simulated impedance and the device’s impedance.

### 3.5 Validation of Directional Stimulation

The optimal simulation parameters identified for omnidirectional stimulation (Dirichlet source, IPG ground) were cross validated against the directional stimulation dataset. The model accurately captured the asymmetric field distribution, yielding a SMAPE of 6.3% at the level of the active segmented contact (Level 2). Fig. 8 illustrates the strong spatial correlation between the simulated and experimental directional fields.

**Figure 8.**
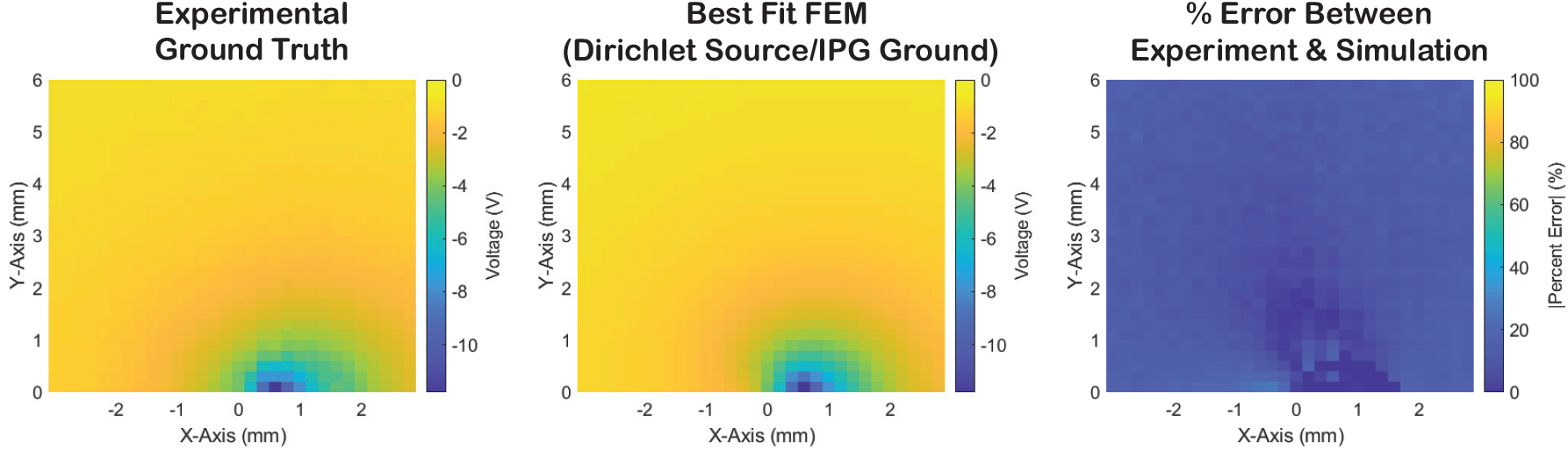
Cross-validation with directional stimulation. The optimal simulation parameters (Dirichlet source, IPG ground) were applied to a directional lead configuration. The simulation accurately captures the asymmetric voltage field (SMAPE = 6.3%), validating the modeling approach for segmented leads.

## 4. Discussion

### 4.1 Experimental Validation of FEM Parameters

In this study, we established an experimental ground truth to resolve a persistent ambiguity in DBS modeling: the selection of appropriate boundary conditions for directional leads. By comparing sub-millimeter voltage measurements against six distinct FEM configurations, we identified that a Dirichlet source with a Dirichlet ground on the IPG surface yields the highest fidelity to physical reality.

A counter-intuitive finding of this work is that Dirichlet boundary conditions outperformed Neumann conditions, even though the clinical IPG operates as a current source in the sense that the total delivered current is strictly controlled. This discrepancy arises from a fundamental conflict in standard FEM solvers between how source constraints are implemented and how real conductive materials behave. Physically, the highly conductive platinum–iridium contacts must remain an equipotential surface. In vivo, although the IPG regulates the total injected current, that current redistributes non-uniformly across the conductor surface in order to satisfy the equipotential condition imposed by the material properties. However, standard Neumann boundary conditions in many FEM solvers enforce a uniform current flux density (*J*) across the boundary. To satisfy this artificial uniform flux constraint, the solver is forced to vary the potential across the contact face, violating the physical behavior of a highly conductive electrode. By instead prioritizing the equipotential constraint (via a voltage boundary condition derived from *V* = *I* × *Z*), the simulation preserves the physically realistic non-uniform current density distribution (including edge effects) that drives neural activation, whereas the standard current boundary condition artificially smooths the electric field.

### 4.2 Clinical Implications: The Risk of VTA Overestimation

The widespread adoption of automated optimization algorithms relies on the accuracy of “pre-computed field libraries”. Our results indicate that unvalidated choices in generating these libraries can introduce systemic bias into clinical predictions.

Most notably, simulations utilizing standard Neumann settings with distant grounds overestimated the VTA by approximately 1.7-fold (137.6 mm^3^ vs. 82.2 mm^3^) compared to the experimentally validated model. In a clinical context, a ∼70% error in volume estimation is non-negligible. It could lead an optimization algorithm to falsely predict side effects (e.g., capsular activation) where none exist, or conversely, to underestimate the spread of stimulation required for therapeutic benefit. These findings suggest that existing commercial and open-source toolboxes should be audited to ensure their underlying physics engines utilize equipotential source models rather than uniform current flux assumptions.

From a clinical standpoint, the 67% VTA overestimation observed in this work has direct implications for patient care, as programming decisions often hinge on millimetric trade offs between benefit and side effects in STN and GPi DBS. In practice, a model that overestimates spread may cause automated algorithms to discard otherwise effective settings as “capsular,” whereas underestimation risks protracted programming sessions that escalate amplitude without achieving durable symptom control. These results should also be interpreted in the context of common clinical parameter ranges, including pulse widths of 60–90 *µ*s (and up to 120 *µ*s in some centers) and the frequent use of bipolar or multipolar configurations to manage side effects or power consumption, which may shift the effective activation threshold relative to the 26.7 V/cm^2^ criterion used here. Finally, the homogeneous saline phantom—while optimal for isolating boundary condition effects—does not capture tissue anisotropy and patient specific anatomy, so VTA visualizations should be viewed as probabilistic guides rather than hard boundaries, and validating these modeling recommendations in subject specific head models remains an important next step.

### 4.3 Practical Recommendations for Modeling

Based on these findings, we recommend that researchers constructing DBS finite element models utilize Dirichlet boundary conditions to enforce the equipotential constraint of the electrode, with the voltage magnitude calculated from the specific tissue impedance and target current (*V* = *I*_target_ × *Z*_measured_). Furthermore, the return path should be explicitly defined by modeling the IPG surface as the 0 V ground; if the IPG cannot be included in the model domain (e.g., in truncated head-only models), our impedance-matched results suggest that constraining the ground to the bottommost surface of the volume conductor provides a superior approximation compared to grounding all outer boundaries.

### 4.4 Limitations

Our experimental setup utilized a homogeneous saline solution to isolate the effects of boundary conditions from tissue heterogeneity. Consequently, these results do not capture the anisotropic conductivity of white matter or the inhomogeneity of encapsulation tissue. However, the physics governing the electrode-electrolyte interface (e.g., the equipotential nature of the contact) are fundamental and independent of the surrounding tissue heterogeneity. Therefore, the superiority of the Dirichlet boundary condition observed here is expected to hold in anatomical models, even if the absolute shape of the VTA changes.

## 5 Conclusion

We provide the first experimental validation of electric field boundary conditions for directional DBS leads using a high-precision robotic phantom. Our results demonstrate that while clinical devices act as current sources, standard Neumann (current density) boundary conditions fail to capture the equipotential nature of the electrode-tissue interface, leading to a significant error (1.7-fold for our experimental setup) in predicted VTA. To ensure the validity of predictive clinical models, we recommend that researchers utilize Dirichlet boundary conditions derived from the device’s operating impedance rather than standard current density settings.

## Data availability

Data is available upon request.

## Supplementary data

Supplementary figures and tables are provided.

