## Supplementary Tables & Figs for "Experimental Validation of Finite Element Models for Directional DBS: The Critical Role of Boundary Conditions on VTA Accuracy"

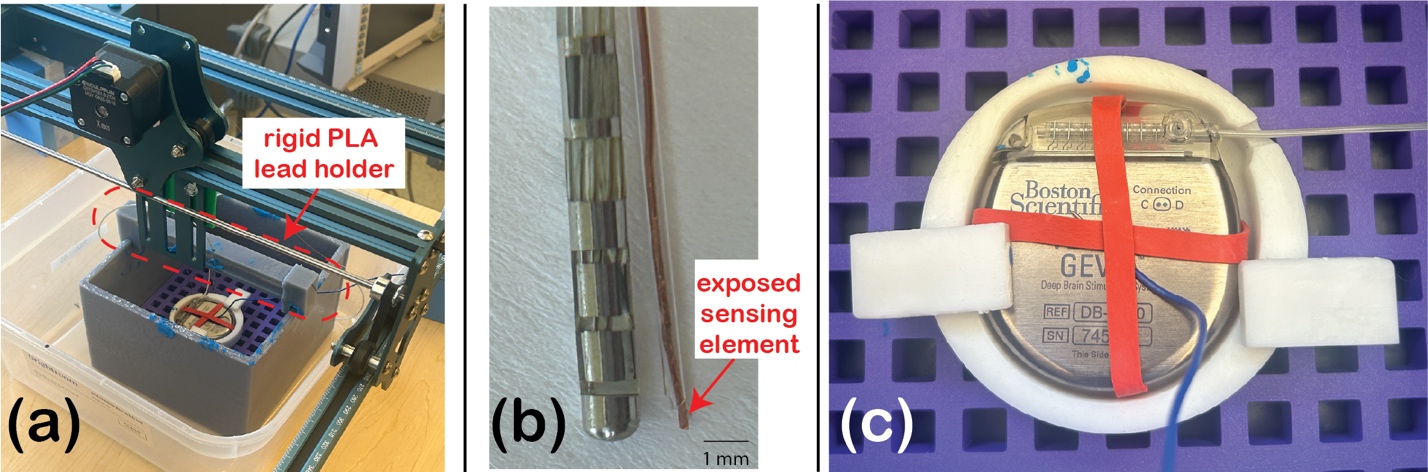

**Supplementary Fig. 1.** (a) The rigid PLA holder is used to keep the DBS lead stable during measurements. (b) The DBS lead and custom-made measurement probe. (c) A top-down view of the IPG and the wire (blue, with exposed tip) we physically connected to the IPG case (the ground).

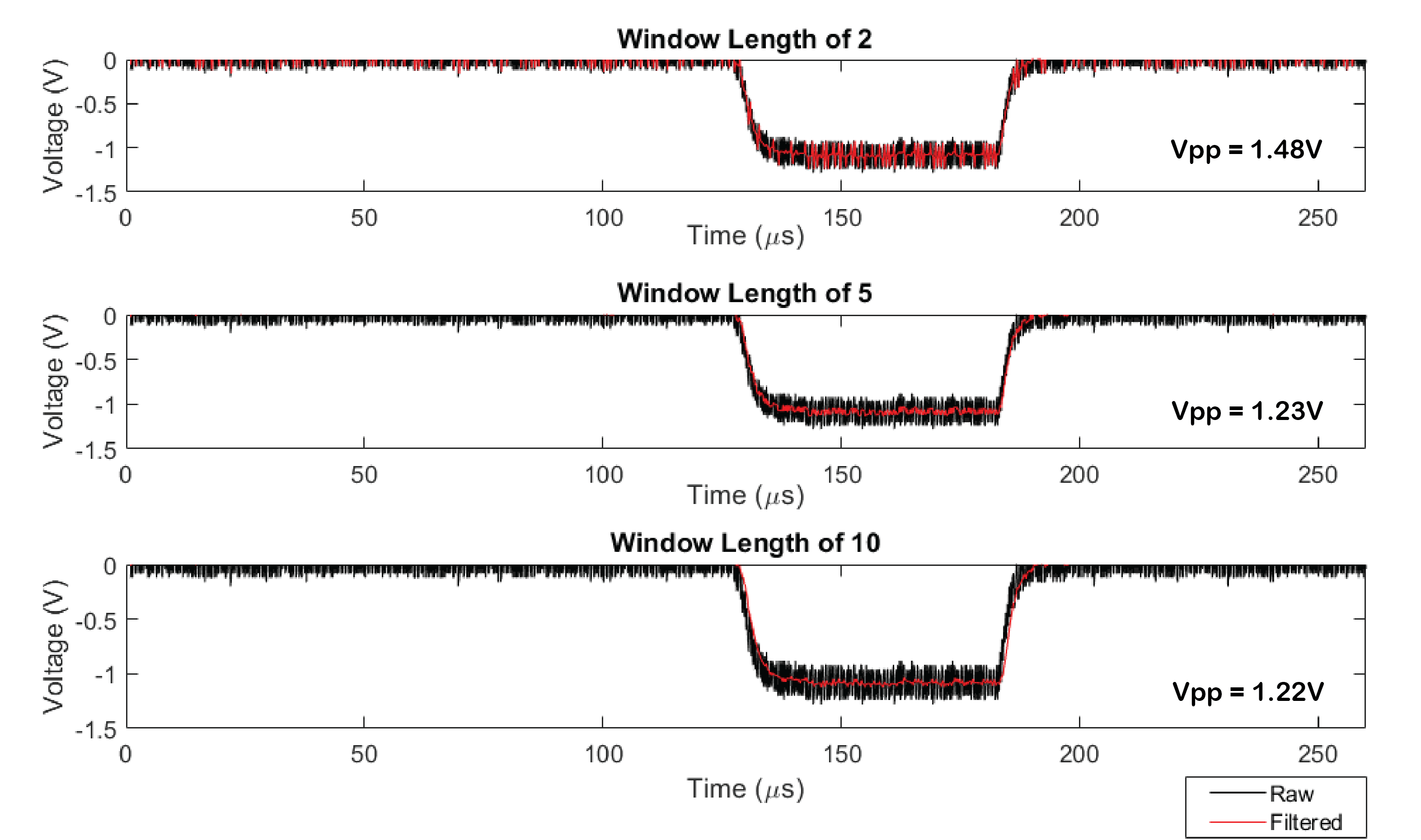

**Supplementary Fig. 2.** Optimization of the moving-average filter kernel size. **(Top)** A narrow window (n=2) fails to sufficiently reject high-frequency noise, resulting in an artificially inflated peak-to-peak voltage (Vpp = 1.48 V) driven by outlier noise spikes. **(Middle)** A window length of 5 samples effectively smooths the noise floor while preserving the true signal amplitude (Vpp = 1.23 V). **(Bottom)** Increasing the window to 10 samples offers negligible additional smoothing benefit (Vpp = 1.22 V). Therefore, a window size of 5 was selected as the optimal trade-off between noise rejection and signal fidelity.

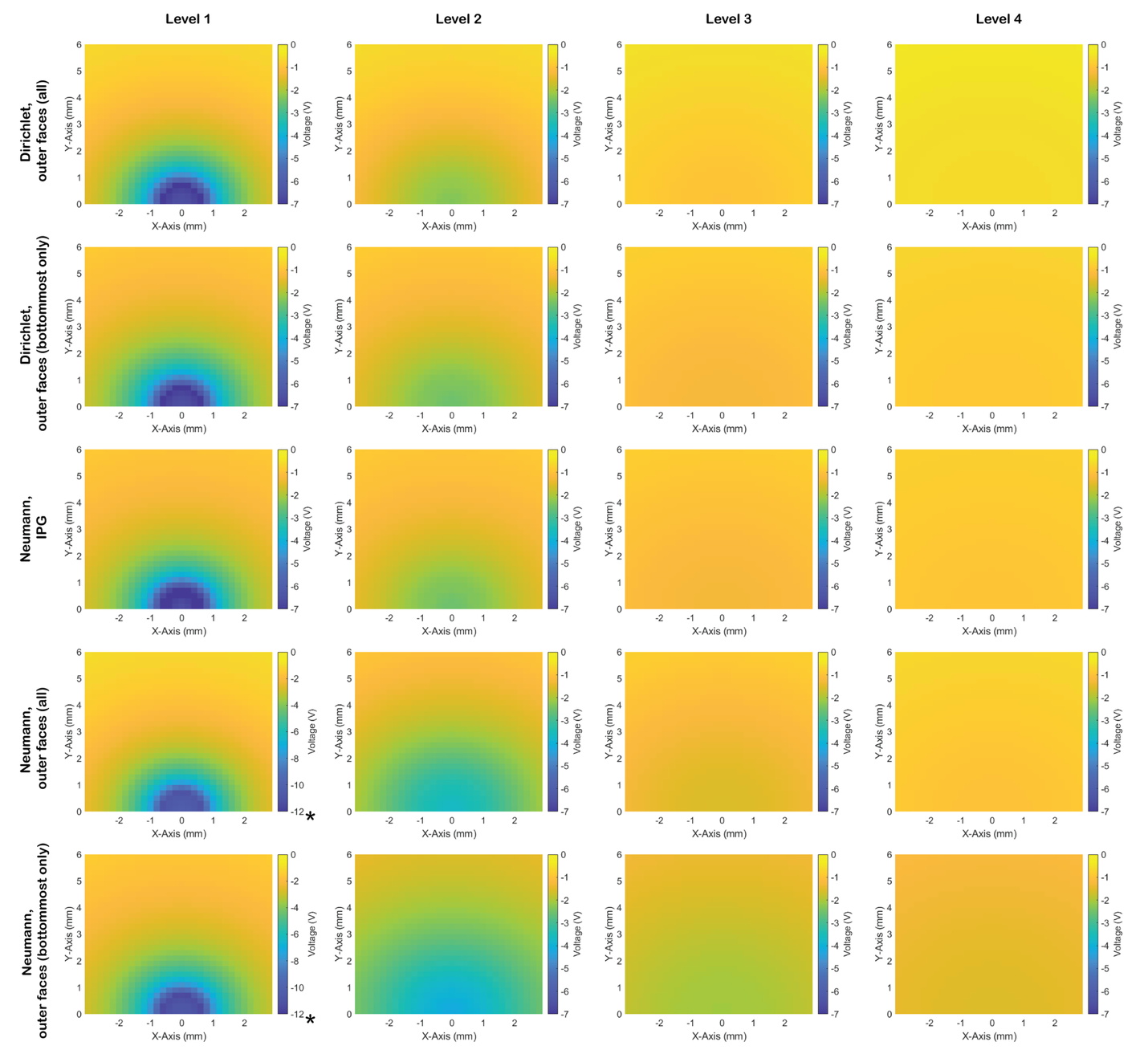

**Supplementary Fig. 3.** The experimental voltage distributions were compared to the simulated voltage distributions at each level of contacts for the simulation setups that were not the best match to the experiment. Note that the color bar range is -7 to 0 V for all distributions except for the two current-controlled simulations on the right for level 1 (denoted with an asterisk).

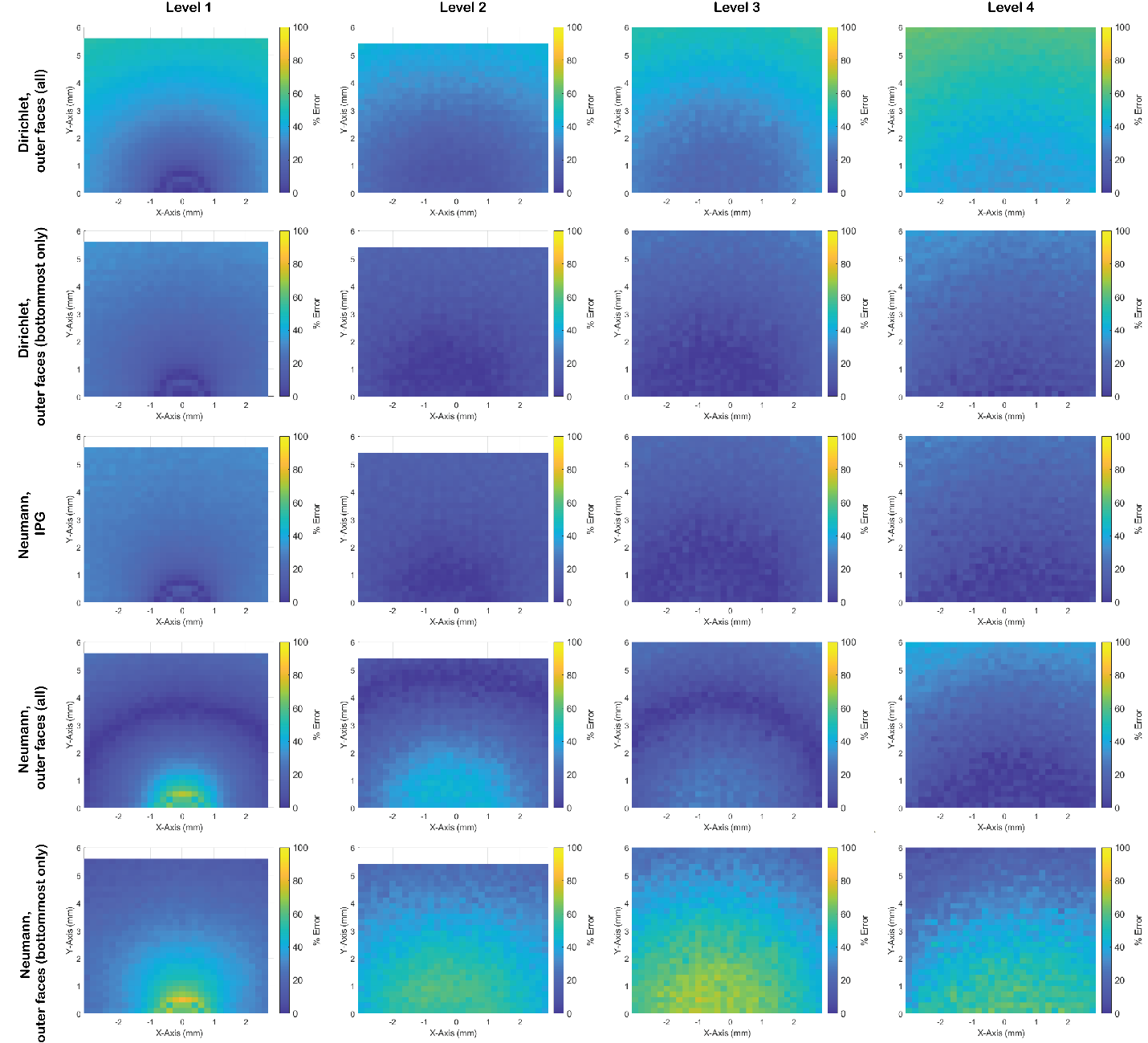

**Supplementary Fig. 4.** Spatial error maps (%) corresponding to the simulation results shown in Supp. Fig. 3.

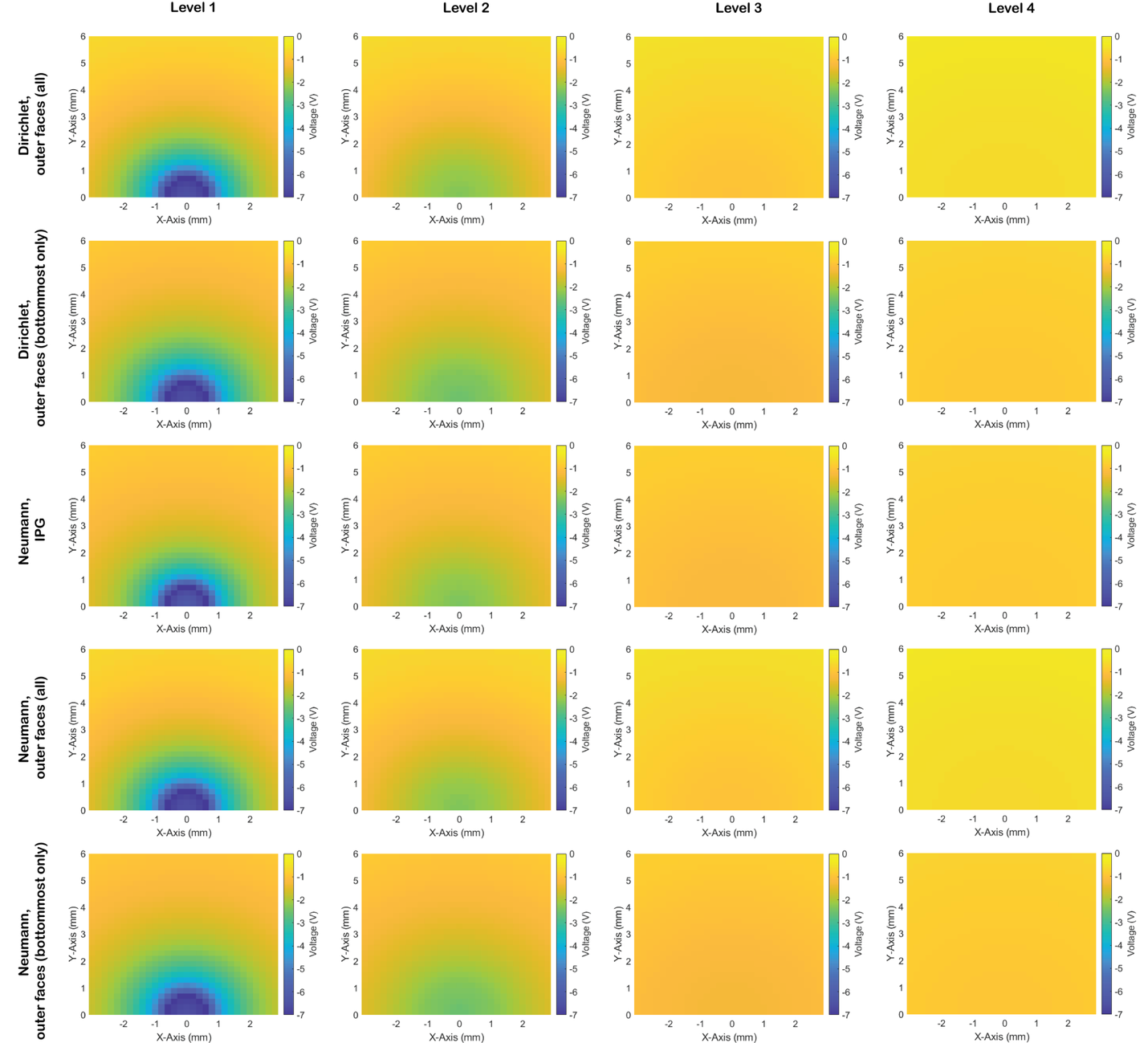

**Supplementary Fig. 5.** The experimental voltage distributions were compared to the simulated voltage distributions (after matching the impedance to the experimental impedance) at each level of contacts for the three simulation setups that were not the best match to the experiment.

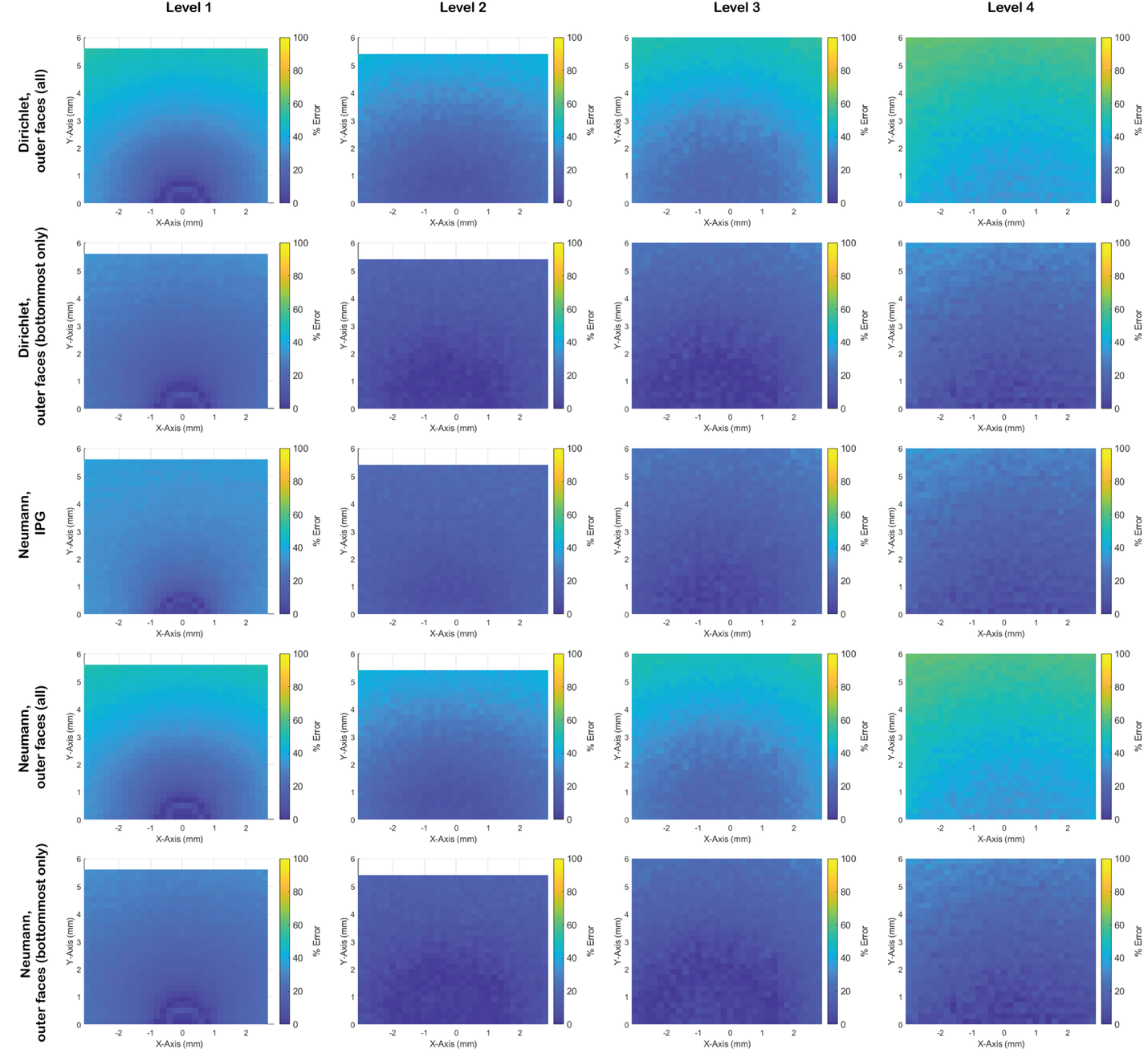

**Supplementary Fig. 6.** Spatial error maps (%) corresponding to the “matching impedance” simulation results shown in Supp. Fig. 5.

**Supplementary Table 1.** Results from comparing the experimentally collected voltage distributions from a cathodic, monopolar setting on contact 8 with the best matching simulation setup. For each level of data collection, the minimum and maximum voltage along the plane, the calculated minimum and maximum absolute percent error, and the SMAPE value are listed.

| Simulation Setup | Level | Simulated  Min (V), Max (V) | Simulation vs. Experiment:  SMAPE (%) |
| --- | --- | --- | --- |
| Dirichlet, IPG  (σ = 0.1 S/m, Z = 1217 Ω) | 1 | -1.16, -0.90 | 4.7 |
|  | 2 | -1.45, -0.96 | 5.3 |
|  | 3 | -2.80, -1.06 | 4.6 |
|  | 4 | -7.06, -1.09 | 4.9 |

**Supplementary Tables 2 – 7:** ANSYS Simulation and mesh details.

- Dirichlet, IPG

|  | Not matching impedance | Matching impedance |
| --- | --- | --- |
| Total # of elements | 3,581,422 | 4,870,753 |
| Percent error | 0.1 | 0.1 |
| Solution Process | Real time = 01:36:15  CPU time = 02:53:02 | Real time = 00:52:46  CPU time = 01:47:43 |
| Number of Passes | 16 out of 20 | 17 out of 20 |

- Neumann, IPG

|  | Not matching impedance | Matching impedance |
| --- | --- | --- |
| Total # of elements | 11,785,530 | 11,785,530 |
| Percent error | 0.1 | 0.1 |
| Solution Process | Real time = 05:59:38  CPU time = 10:04:38 | Real time = 05:33:12  CPU time = 09:47:15 |
| Number of Passes | 18 out of 20 | 18 out of 20 |

- Dirichlet, outer faces (all)

|  | Not matching impedance | Matching impedance |
| --- | --- | --- |
| Total # of elements | 290,052 | 290,052 |
| Percent error | 0.1 | 0.1 |
| Solution Process | Real time = 00:08:14  CPU time = 00:21:25 | Real time = 00:08:02  CPU time = 00:21:28 |
| Number of Passes | 17 out of 20 | 17 out of 20 |

- Neumann, outer faces (all)

|  | Not matching impedance | Matching impedance |
| --- | --- | --- |
| Total # of elements | 1,835,799 | 1,835,799 |
| Percent error | 0.1 | 0.1 |
| Solution Process | Real time = 01:00:52  CPU time = 02:44:53 | Real time = 00:59:54  CPU time = 02:50:22 |
| Number of Passes | 23 out of 26 | 23 out of 26 |

- Dirichlet, outer faces (bottommost only)

|  | Not matching impedance | Matching impedance |
| --- | --- | --- |
| Total # of elements | 213,262 | 213,262 |
| Percent error | 0.1 | 0.1 |
| Solution Process | Real time = 00:5:58  CPU time = 00:15:41 | Real time = 00:05:34  CPU time = 00:14:44 |
| Number of Passes | 16 out of 20 | 16 out of 20 |

- Neumann, outer faces (bottommost only)

|  | Not matching impedance | Matching impedance |
| --- | --- | --- |
| Total # of elements | 1,349,810 | 1,349,810 |
| Percent error | 0.1 | 0.1 |
| Solution Process | Real time = 00:43:00  CPU time = 02:01:42 | Real time = 00:42:20  CPU time = 01:59:13 |
| Number of Passes | 22 out of 26 | 22 out of 26 |
